# Shared internal models for feedforward and feedback control of arm dynamics in non-human primates

**DOI:** 10.1101/2020.04.05.026757

**Authors:** Rodrigo S. Maeda, Rhonda Kersten, J. Andrew Pruszynski

## Abstract

Previous work has shown that humans account for and learn novel properties or the arm’s dynamics, and that such learning causes changes in both the predictive (i.e., feedforward) control of reaching and reflex (i.e., feedback) responses to mechanical perturbations. Here we show that similar observations hold in old-world monkeys (macaca fascicularis). Two monkeys were trained to use an exoskeleton to perform a single-joint elbow reaching and to respond to mechanical perturbations that created pure elbow motion. Both of these tasks engaged robust shoulder muscle activity as required to account for the torques that typically arise at the shoulder when the forearm rotates around the elbow joint (i.e., intersegmental dynamics). We altered these intersegmental arm dynamics by having the monkeys generate the same elbow movements with the shoulder joint either free to rotate, as normal, or fixed by the robotic manipulandum, which eliminates the shoulder torques caused by forearm rotation. After fixing the shoulder joint, we found a systematic reduction in shoulder muscle activity. In addition, after releasing the shoulder joint again, we found evidence of kinematic aftereffects (i.e., reach errors) in the direction predicted if failing to compensate for normal arm dynamics. We also tested whether such learning transfers to feedback responses evoked by mechanical perturbations and found a reduction in shoulder feedback responses, as appropriate for these altered arm intersegmental dynamics. Demonstrating this learning and transfer in non-human primates will allow the investigation of the neural mechanisms involved in feedforward and feedback control of the arm’s dynamics.

## Introduction

Monkeys, like humans, perform arm movements that require coordinating multiple joints. One complexity of such movements are the mechanical interactions that arise across moving limb segments, as movements around one joint cause rotational forces at other joints (Hollerbach & Flash, 1982; Graham *et al*., 2003). Many previous studies have demonstrated that the nervous system develops an internal representation of such arm dynamics that can be used to generate predictive (i.e., feedforward) muscle activity during self-initiated reaching (Gribble & Ostry, 1999; Debicki & Gribble, 2005; Gritsenko *et al*., 2011; Maeda *etal*. 2017, 2018; Maeda, Gribble, *etal*. 2020; Maeda, Zdybal, *etal*. 2020a, 2020b) and also reflex (i.e., feedback) responses to mechanical perturbations, in a way that accounts for the arm dynamics (Lacquaniti & Soechting, 1984, 1986a, 1986b; Soechting & Lacquaniti, 1988; Kurtzer, Pruszynski, *et al*., 2006; Kurtzer *et al*., 2008, 2009, 2014, 2016; Pruszynski *et al*., 2011; Crevecoeur *et al*., 2012; Maeda *et al*., 2017, 2018; Kurtzer, 2019; Maeda, Gribble, *et al*., 2020). For instance, when generating single-joint elbow movements and when responding to mechanical perturbations that create pure elbow motion, the nervous system generates predictive shoulder muscle activity and robust shoulder feedback responses to counter the underlying torques that arise at the shoulder joint because of forearm rotation about the elbow joint (Gribble & Ostry, 1999; Kurtzer *et al*., 2008; Maeda *et al*., 2017, 2018; Maeda, Gribble, *et al*., 2020). This similarity between feedforward and feedback control reflects, in part, the shared neural circuits between feedforward and feedback control at the level of the spinal cord, brainstem and cerebral cortex (Pruszynski & Scott, 2012; Kurtzer, 2014; Scott, 2016).

One way of determining how the nervous system represents arm dynamics is by altering the normal mapping between joint torques and joint motion. For example, we have recently shown in humans that generating pure elbow movements with the shoulder joint fixed, causes people to reduce shoulder muscle contraction during self-initiated reaching and that such learning transfers to feedback responses (Maeda *et al*., 2018; Maeda, Zdybal, *et al*., 2020a). Such learning and transfer is appropriate and efficient because mechanically fixing the shoulder joint eliminates the interaction torques that arise at the shoulder when the forearm rotates and thus removes the need to recruit shoulder muscles.

Here we address the same question in old-world monkeys (macaca fascicularis). Multi-joint arm movements in monkeys account for arm intersegmental dynamics (Graham *et al*., 2003) and monkeys generate predictive underlying muscle activity (Scott, 1997; Gribble & Scott, 2002; Kurtzer, Herter, *et al*., 2006) but it is unknown whether and how they learn new muscle activity patterns to adapt to novel arm dynamics. It is also unknown whether such learning transfers to feedback responses in monkeys. It is important to emphasize that neither the learning or transfer phenomena need to exist because, in our paradigm, the pre-fixation pattern of muscle activity would continue to yield successful outcomes with the shoulder fixed for both self-initiated reaching and when responding to mechanical perturbations. Therefore, establishing these phenomena in monkeys at the level of muscle activity is a critical starting point for detailing the underlying neuronal circuits and mechanisms that make this type of learning and transfer possible.

We trained two monkeys to sit and use a robotic exoskeleton to generate self-initiated elbow reaching rotations and to compensate for mechanical perturbations that created pure elbow motion. As in our human experiments, we altered the normal mapping between joint torque to joint motion by fixing the shoulder joint of the robot. First, we tested whether monkeys adapt to this altered arm dynamics during reaching, and we found that, like humans, monkeys show a reduction in shoulder muscle activity over trials. Second, we tested whether this new feedforward coordination pattern is expressed in feedback responses. As in humans, we found a reduction in shoulder reflex responses when a mechanical perturbation is occasionally introduced after learning. Taken together, these results suggest that, like in humans, feedforward and feedback control in macaque monkeys share an internal model of the arm’s dynamics.

## Materials and Methods

### Animals and apparatus

The Institutional Animal Care and Use Committee at Western University approved all the procedures described below. Two male cynomolgus monkeys (macaca fascicularis, Monkeys A and O, ∼9 and 13 kgs, respectively) were trained to perform a range of upper-limb tasks while seated in a robotic exoskeleton (NHP KINARM, Kingston, Ontario).

As described previously (Scott, 1999; Pruszynski *et al*., 2011, 2014), this robotic device permits flexion and extension movements of the shoulder and elbow joints in the horizontal plane, and can independently apply torque at both of these joints. Target and hand feedback were projected from an LCD monitor onto a semi-silvered mirror, in the horizontal plane of the task. Direct vision of the monkey’s arm was blocked with a physical barrier, and the ambient lights in the experimental room were turned off for the duration of data collection.

The two segments of the exoskeleton robot (upper arm and forearm) were adjusted to fit each monkey’s arm. Monkey A used their left arm while monkey O used their right arm (Figure 1A-B). The robot was then calibrated so that the projected hand cursor was aligned with the tip of each monkey’s index finger. Calibration was saved and used across training and testing sessions. At the beginning of each session, an experimenter gently strapped each monkey’s arm to the two links of the robot, to ensure a tight coupling, and certified that the calibrated hand cursor matched the actual index finger for each animal.

**Figure 1:**
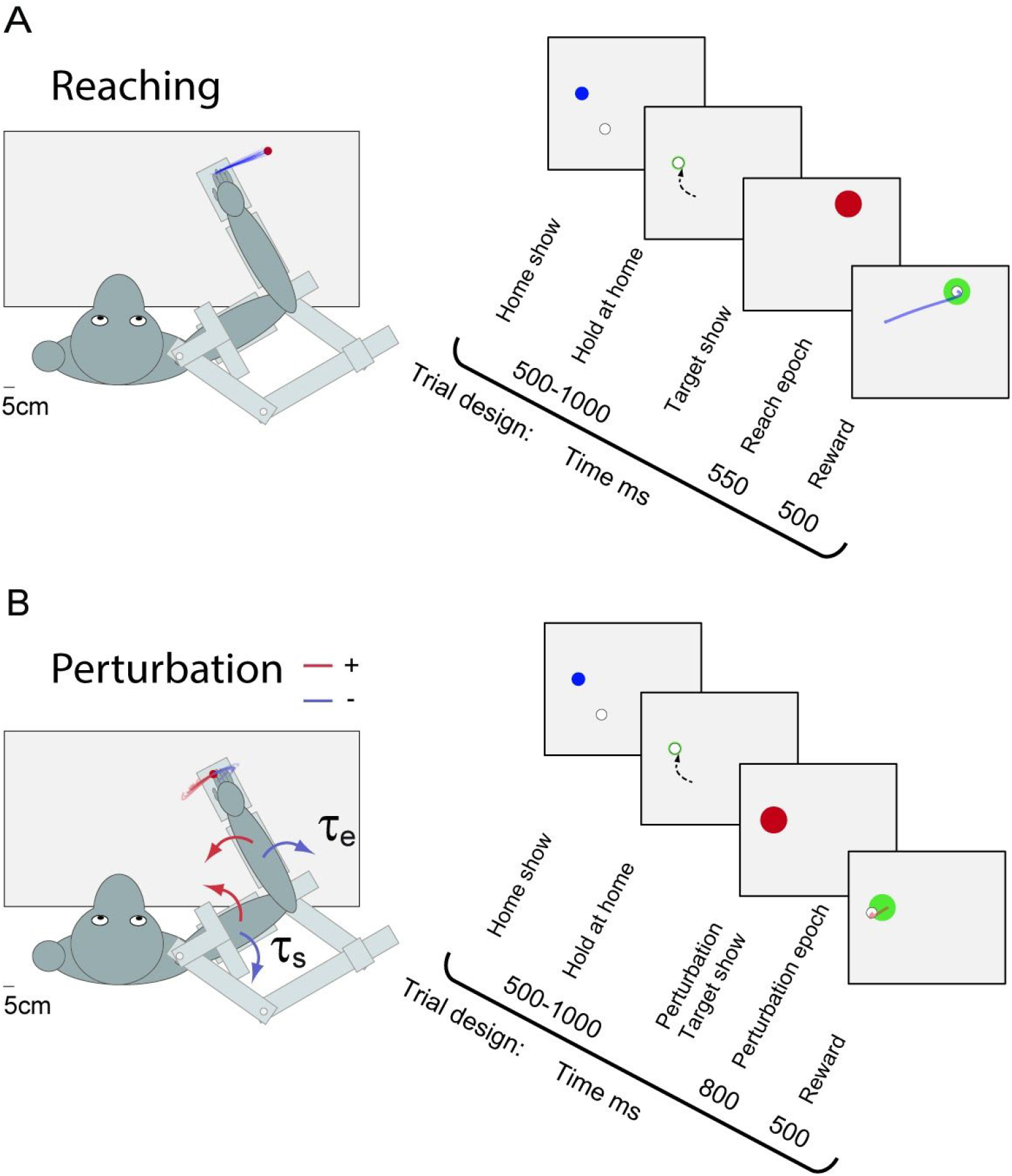
Task and trial design. **A**, Reaching task set-up with individual hand kinematic traces of elbow extension movements from one exemplar session with the shoulder joint free to move (i.e., baseline trials). At the beginning of a reaching trial, a home target was displayed (blue circle, 0.5 cm diameter). Monkeys were trained to move their hand cursor into this home target, and when they did so the home target turned green. After maintaining their hand cursor at this home location for a random period (500 –1000 ms, uniform distribution), a goal target (red circle: 0.8 cm diameter) was presented in a location that could be reached with a 20° of pure elbow extension movement. The monkey received water reward and visual feedback (the goal target turned green) if it transported its hand to the goal target within 550 ms and remained in it for an additional 500 ms. **B**, Perturbation task set-up with individual hand kinematic traces of flexion (red) and extension (blue) shoulder and elbow torque perturbations. In perturbation trials, after maintaining the cursor in the home target for a randomized duration (500 –1000 ms, uniform distribution), a torque pulse (i.e., perturbation) of 100 ms duration was applied simultaneously to the shoulder and elbow joints (0.48 and 0.35 N m, respectively), which displaced the monkey’s hand outside the home target (bottom left row). At the same time as the perturbation, a goal target (red circle: 0.8 cm diameter) was overlaid at the home target location. The monkey received a water reward and visual feedback (the goal target turned green) if it returned its hand to the goal target within 800 ms of perturbation onset and remained in it for an additional 500 ms.

### Reaching and posture perturbation tasks

Monkeys performed a version of the reaching and posture perturbation tasks described and performed by human participants in Maeda et al. (2018). Briefly, each monkey was trained to perform 20° of elbow flexion and extension movements or to counter mechanical perturbations applied by the robot that caused pure elbow motion. Animals did so with the shoulder joint either free to move or with the shoulder joint fixed. Shoulder fixation was achieved by inserting a physical pin into the robot linkage that permits shoulder motion.

At the beginning of a trial, a home target was displayed (blue circle, 0.5 cm diameter). The home target position corresponded to each monkey’s hand cursor position (white circle, 0.4 cm diameter), when their shoulder and elbow joint was positioned at 35° and 60°, respectively (external angles; Figure 1A, left column). Monkeys were trained to move their hand cursor into this home target, and when they did so the home target turned green. After maintaining their hand cursor at this home location for a random period (500 –1000 ms, uniform distribution), a goal target (red circle: 0.8 cm diameter) was presented in a location that could be reached with a 20° of pure elbow flexion movement. The monkey received water reward and visual feedback (the goal target turned green) if it transported its hand to the goal target within 550 ms and remained in it for an additional 500 ms (Figure 1A, right column). For the trial to be considered a correct trial, the hand cursor was required to end inside the target. After an interval period of 1000 ms, the goal target became a new home target (blue circle, 0.5 cm diameter) and the same procedure was repeated for an extension movement.

Perturbation trials occurred in 15% of all trials. In these trials, after maintaining the cursor in the home target for a randomized duration (500 –1000 ms, uniform distribution), a torque pulse (i.e., perturbation) of 100 ms duration was applied simultaneously to the shoulder and elbow joints (0.48 and 0.35 N m, respectively), which displaced the monkey’s hand outside the home target (Figure 1B, left column). We chose this combination of shoulder and elbow loads to minimize shoulder motion (see Kurtzer *etal*., 2008, 2009, 2014; Pruszynski *etal*., 2011; Maeda *etal*., 2017, 2018; Kurtzer, 2019; Maeda, Gribble, *etal*., 2020; Maeda, Zdybal, *etal*., 2020a). At the same time as the perturbation, a goal target (red circle: 0.8 cm diameter) was overlaid at the home target location. The monkey received a water reward if it returned its hand to the goal target within 800 ms of perturbation onset and remained in it for an additional 500 ms (Figure 1B, right column). For the trial to be considered a correct trial, the hand cursor was required to end inside the target. If the monkey returned to the goal target within this timing window, the target circle changed from red to green, otherwise the target circle remained red and no reward was provided. Note that perturbations could occur at either the more flexed or extended home target positions.

Monkeys performed blocks of trials, each consisting of 20 flexion and extension reaching trials and 4 perturbation trials. The order of reaching and perturbation trials was randomized within a block of trials. Recording sessions consisted of 1-2 blocks of trials with the shoulder joint free to move (baseline phase), 4-8 blocks of trials with the shoulder joint of the robot fixed (adaptation phase), and 1-2 blocks with the shoulder joint free to rotate again (post-adaptation phase). The precise number of blocks was determined by the experimenter based on animal behavior on that day (see Table 1).

**Table 1.**
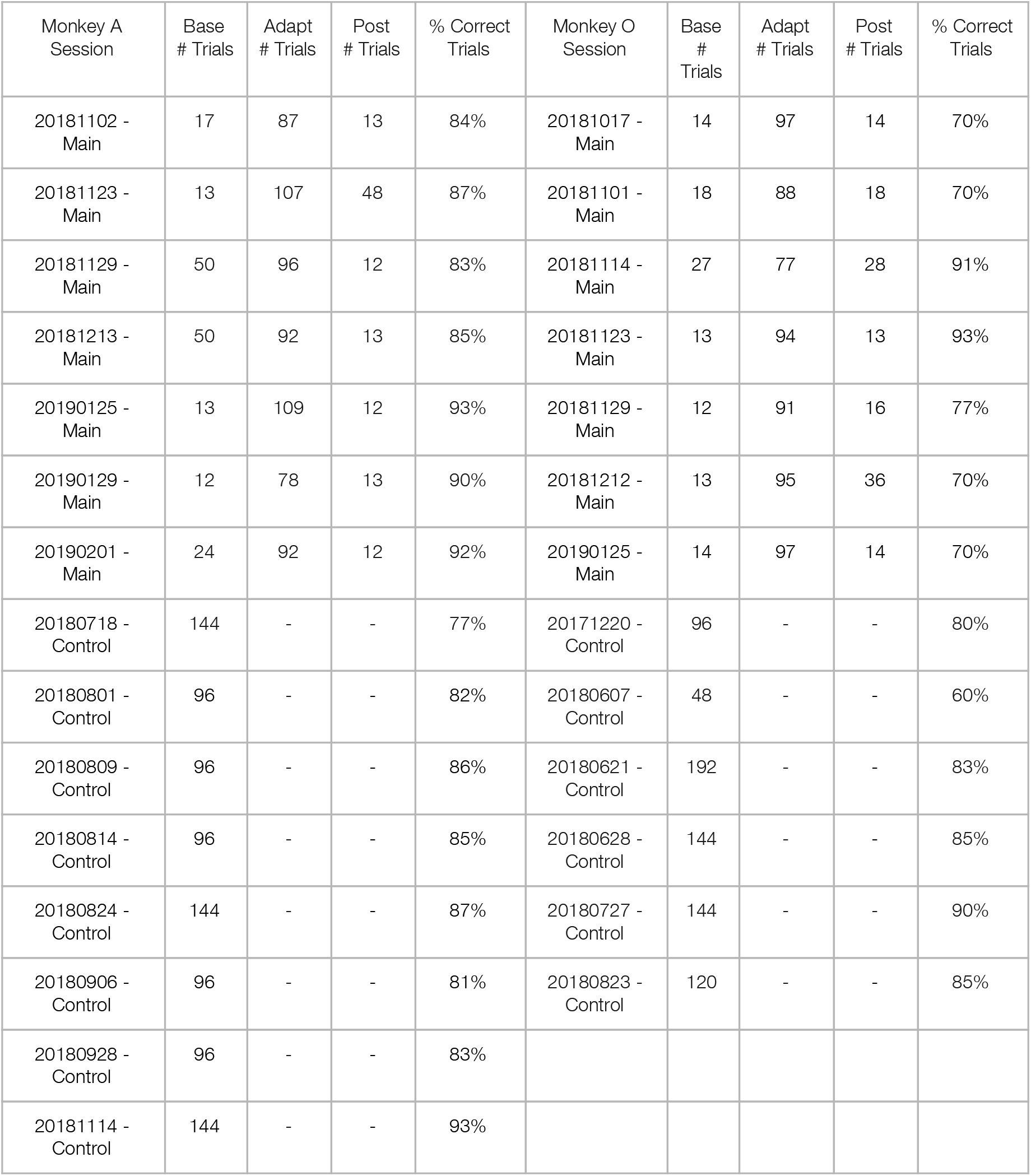
Number of extension trials and % of correct trials in each session for monkeys A and O.

Five to ten additional sessions were recorded from each animal using the same version of this protocol but without locking the shoulder joint. This served as a control to rule out changes caused by extensive movement repetitions within a session rather than the shoulder fixation manipulation (see Table 1).

Experimental sessions lasted approximately 2-2.5 hours. Rest breaks were given between blocks of trials. Before data collection, monkey A completed ∼1 year and monkey O completed ∼2 years of positive reinforcement training in this task. Each animal completed these training sessions with all the joints free to rotate and never experienced shoulder fixation over the course of training until the experimental sessions being reported here.

### Muscle and kinematic recordings

We acquired bipolar electromyographic (EMG) activity, using intramuscular fine-wire electrodes. EMG electrodes were made in-house and consisted of Teflon-coated 3 or 7 -strand stainless steel wire with 50 to 100 mm length (A-M Systems, Sequim, WA), threaded into a 30-mm, 27-gauge cannula (Kendall Monoject, Mansfield, MA), and hooked to anchor to the muscle tissue. Electrodes were spaced ∼8-10mm apart and aligned to the muscle fibers.

We focused on five upper limb muscles [pectoralis major clavicular head (PEC), shoulder flexor; posterior deltoid (PD), shoulder extensor; biceps brachii long head (BB), shoulder and elbow flexor, brachioradialis (BR), elbow flexor; triceps brachii lateral head (TR), elbow extensor]. These muscles were chosen because they are easily accessible and active during flexion and extension movements at the shoulder and elbow joints (Kurtzer, Herter, *et al*., 2006; Kurtzer, Pruszynski, *et al*., 2006). A reference electrode was inserted subcutaneously on the animal’s back. To decrease stress, animals were sometimes given a mild sedative prior to the EMG placement. The sedation used was a combination of medetomidine (0.01-0.02mg/kg) and midazolam (0.04-0.05mg/kg). The medications were given by intramuscular injection in the home cage. Immediately following the injection animals were moved into the KINARM chair and taken to the procedure room where the lights were dimmed to reduce stimulation. Once the animals were relaxed (usually 10-15min post injection) insertion of the EMG wires into the above mentioned muscle locations started. Once all of the wires were placed, the Medetomidine portion of the sedation was reversed using Atipamezole intramuscularly (volume same as medetomidine administered in that session).

Muscle activity was recorded at 2,000 Hz, bandpass filtered (25–500 Hz, sixth order Butterworth) and full-wave rectified before analysis (Pruszynski *et al*., 2014). Kinematic data and applied torques were sampled at 1,000 Hz, directly from the KINARM device, and then low-pass filtered before analysis (12 Hz, 2-pass, fourth-order Butterworth).

### Data and statistical analysis

Data from reaching trials were aligned on movement onset and data from perturbation trials were aligned on perturbation onset. Movement onset was defined as 10% of peak angular velocity of the elbow joint. Only rewarded trials were selected for analysis. Only those muscle recordings sessions that yielded clear phasic responses to either movement or mechanical perturbation, and minimal motion artifact, were analyzed. Specifically, we focused our key analyses on PD and TR muscle activity because the other muscles did not provide reliable reliable and high quality signals across sessions and across feedforward and feedback experimental conditions. EMG during reaching was normalized to the average activity in a fixed time window (−100 to +100 ms, relative to movement onset, capturing the agonist burst of EMG activity) in baseline trials, such that a value of 1 represents a given mean activity when a given muscle is acting as agonist. EMG during perturbation was normalized to the average activity in the voluntary response epoch (+100 to +150 ms, relative to perturbation onset, known to capture the agonist burst of EMG activity) in baseline trials, such that a value of 1 represents a given mean activity in the voluntary response when a given muscle is acting as agonist.

Our main interest here was to investigate whether monkey’s shoulder (specifically, PD) muscle activity during self-initiated reaching is reduced following shoulder fixation. To answer this question, we calculated the mean amplitude of phasic PD muscle activity in reaching trials across a fixed time window (−100 to +100 ms, relative to movement onset and chosen to capture the agonist burst of EMG activity). We then tracked whether and how this measurement of muscle activity changed over time and across different phases of the protocol with the shoulder joint free and fixed (Debicki & Gribble, 2005; *etal*., 2018; Maeda, Gribble, *et al*., 2020; Maeda, Zdybal, *et al*., 2020a, 2020b).

We also quantified kinematic aftereffects of reaching movements after adapting reaching with shoulder fixation. To do so, we calculated the hand path errors relative to the center of the target at 80% of the movement between movement onset and offset (also defined at 10% of peak angular velocity of the elbow joint). This window was chosen to select the kinematic traces before any feedback corrections (Maeda *et al*., 2018; Maeda, Gribble, *et al*., 2020; Maeda, Zdybal, *et al*., 2020a).

Our second goal was to investigate whether feedback responses in shoulder (specifically, PD) muscle activity also adapt to the novel arm dynamics learned during self-initiated reaching with shoulder fixation. To answer this question, we binned the PD EMG of perturbation trials into previously defined epochs (Pruszynski *et al*., 2008; Maeda *et al*., 2018). This included a pre-perturbation epoch (PRE; 50 – 0 ms relative to perturbation onset), the short-latency stretch response (20–50 ms), the long-latency stretch response (50 – 100 ms), and the voluntary response (100 –150 ms).

All data processing and analyses were performed using MATLAB (r2019a, MathWorks). We performed different statistical tests (e.g., repeated-measures ANOVA with Tukey tests for multiple comparisons, t test), when appropriate for each experimental question. Statistical analyses were performed using R (v3.2.1, R Foundation for Statistical Computing). Details of these procedures are provided in Results. Experimental results were considered statistically reliable if the corrected p-value was <0.05.

## Results

### Compensating for arm dynamics during reaching and perturbation trials

Both monkeys were successful at achieving the imposed speed and accuracy constraints of the tasks in the testing sessions (average success rate 85% for monkey A and 78% for monkey O, see Table 1).

Even with the shoulder free to move (i.e., baseline trials), monkeys moved from the home target to the goal target with minimal motion at the shoulder joint and with substantial motion at the elbow joint (Figure 2A-B), similar to what humans do when presented with similar targets and without instruction about how to move (Maeda *et al*., 2017, 2018). Despite minimal shoulder rotation, we found substantial shoulder extensor muscle activity before movement onset (10% of peak elbow velocity; see Materials and Methods), as required to compensate for the torques that arise at the shoulder when the forearm rotates (Gribble & Ostry, 1999; Maeda *et al*., 2017, 2018; Maeda, Gribble, *et al*., 2020; Maeda, Zdybal, *et al*., 2020a; Figure 2C-D).

**Figure 2:**
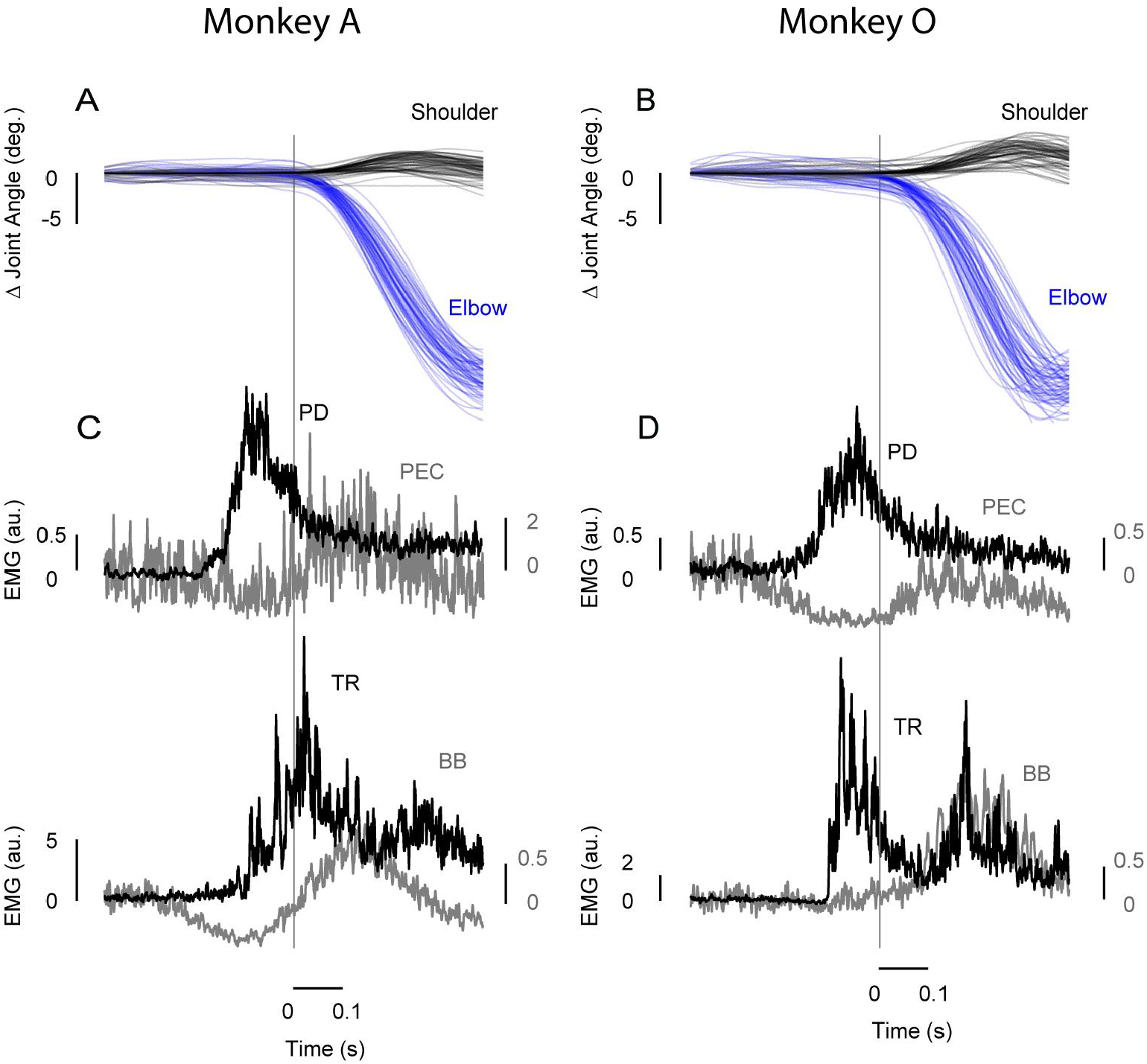
Accounting for arm dynamics during feedback responses to perturbation. **A-B**, Kinematics of the shoulder (black) and elbow (blue) joints for elbow extension movements for monkeys A and O, respectively. Data are aligned on movement onset. Data is shown for individual trials. **C-D**, Average agonist (black traces: PD, top row and TR, bottom row) and antagonist (gray traces: PEC, top row and BB, bottom row) muscle activity of the movements in **A-B** for monkeys A and O, respectively. EMG data are normalized as described in the Methods. Data are aligned on movement onset. **A-D**, All data from the same exemplar session.

In selected trials, we occasionally applied mechanical perturbations to the arm while the monkeys maintained their hand in the home target. The mechanical perturbations consisted of 100ms torque pulses applied simultaneously to the shoulder and elbow joints with magnitudes chosen to elicit minimal shoulder motion but different amounts of elbow motion (Figure 3A-B and inset). Consistent with previous work, we found that these mechanical perturbations elicited substantial shoulder muscle activity in the long-latency epoch (50-100 ms post-perturbation; Figure 3C-D), as appropriate for countering the imposed joint torques (Kurtzer *etal*., 2008, 2009, 2014; Pruszynski *etal*., 2011; Maeda *etal*., 2017, 2018; Kurtzer, 2019; Maeda, Gribble, *etal*., 2020; Maeda, Zdybal, *etal*., 2020a).

**Figure 3:**
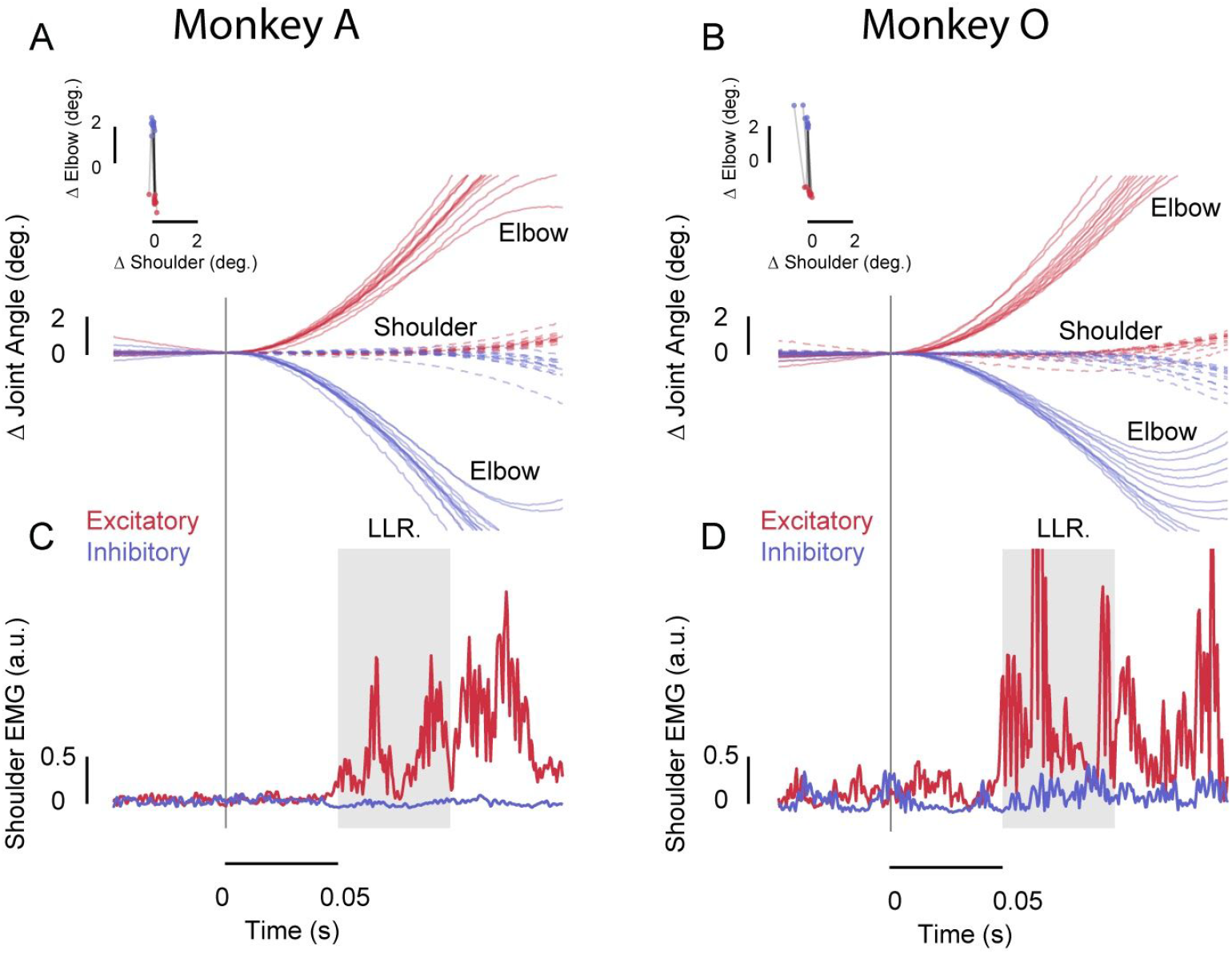
Accounting for arm dynamics during feedback responses to perturbation. **A-B**, Kinematic traces (top), shoulder (dashed) and elbow (solid) joint kinematics during perturbation trials for monkeys A and O, respectively. Data are aligned on perturbation onset. Data is shown for individual trials. Red and blue traces are from the shoulder/elbow flexor torque and shoulder/elbow extensor torque perturbation conditions, respectively. Inset shows the amount of shoulder and elbow displacement 50 ms post-perturbation (data are shown for individual trials in this session). **C-D**, Average shoulder PD muscle activity following mechanical perturbations with the shoulder joint free to move (i.e., baseline trials). Data are aligned on perturbation onset. EMG data are normalized as described in the Methods. Grey horizontal shaded area indicates the Long-Latency Reflex epoch (LLR). **A-D**, All data from the same exemplar session.

### Learning single-joint elbow reaching with shoulder fixation

We first asked whether monkeys adapt reaching movements to the imposed novel arm dynamics with shoulder fixation, a manipulation that eliminates the torques that arise at the shoulder and that need to be compensated by muscles at the shoulder joint with motion of the forearm. Note that monkeys were not exposed to shoulder fixation in the training sessions, but when it was introduced in the testing sessions, fixation of the shoulder joint did not alter task performance, and monkeys continued to demonstrate success rates of 85% for monkey A and 78% for monkey O (see Table 1).

Figure 4A, E illustrates, for both monkeys, the mean shoulder extensor muscle activity in a fixed time window (−100 to +100 ms, relative to movement onset; see Materials and Methods), across elbow extension trials before (i.e., baseline phase), during (i.e., adaptation phase) and after (i.e., adaptation phase) the shoulder was physically fixed at the robotic manipulandum. The amplitude of shoulder muscle activity was reduced over the course of adaptation trials with the shoulder joint fixed and quickly returned to baseline levels after releasing the shoulder joint again (i.e., post-adaptation phase).

**Figure 4:**
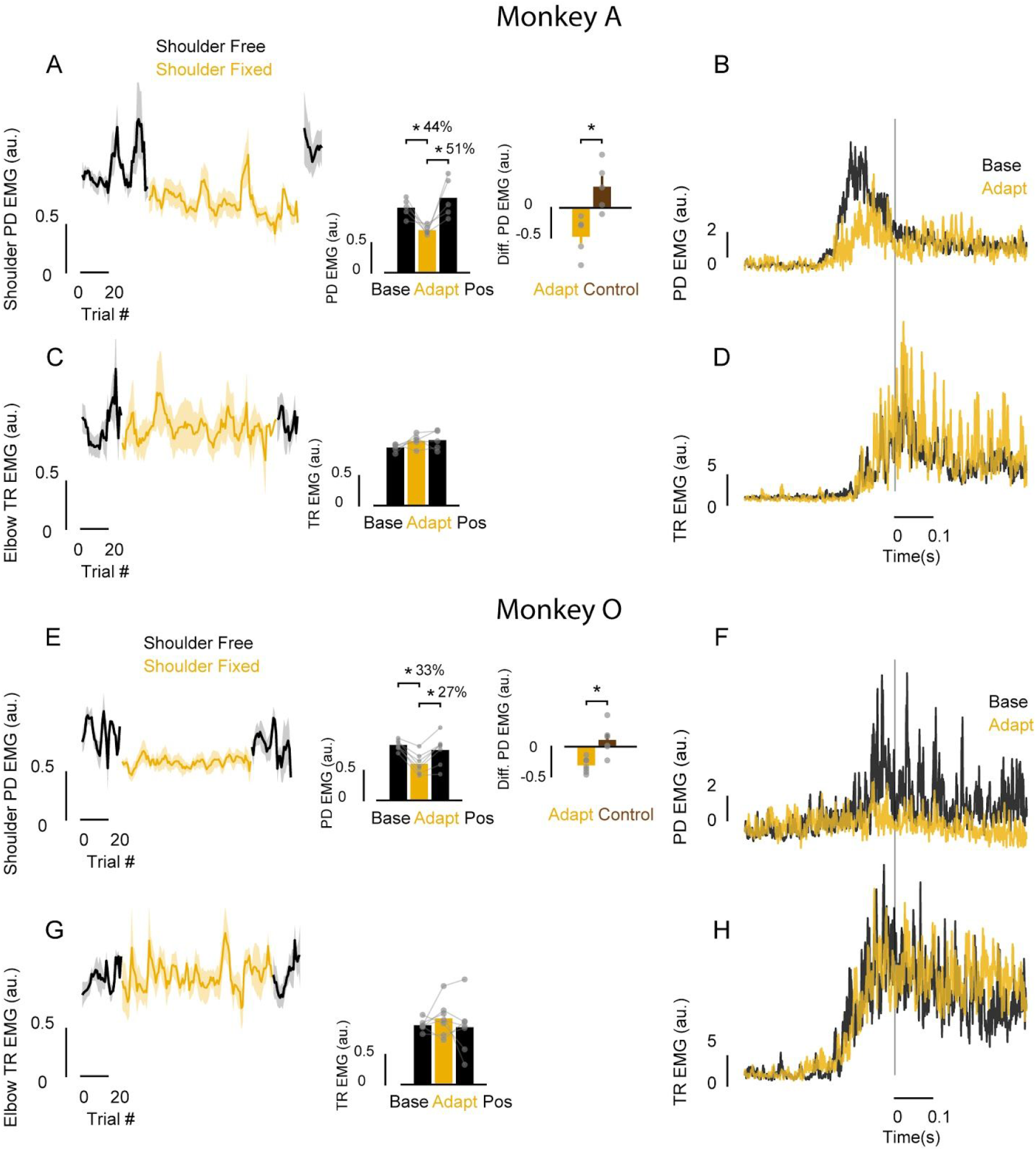
Learning novel arm dynamics during elbow reaching with shoulder fixation. **A**, Normalized PD muscle activity in a fixed time window (−100 to +100 ms relative to movement onset), averaged across sessions and smoothed over trials (i.e., moving average of 5 trials) with the shoulder free (black) and fixed (yellow). Shaded error areas represent standard error of the mean (SE). Left inset bar graph shows the average normalized PD muscle activity (−100 to +100 ms relative to movement onset) associated with these trials late in the baseline, adaptation and post-adaptation phases. Each dot represents data from a single session. Asterisks indicate reliable effects (p < 0.05; see main text). Right inset bar graph shows the difference in shoulder PD muscle activity (late of adaptation - baseline phases) for sessions in which the shoulder was fixed (adapt) and sessions in which the shoulder was never fixed (control). **B**, Time series of normalized PD muscle activity in one exemplar session averaged over the last 20 baseline and adaptation trials. Data are aligned on movement onset. **C-D**, Data for TR muscle are shown using the same format as **A-B. E-H**, Data for monkey O are shown using the same format as for monkey A in **A-D**.

A one-way ANOVA comparing shoulder extensor muscle activity late (last 20 trials) in the baseline, adaptation and post-adaptation phases indeed revealed a reliable effect of phase on shoulder muscle activity across sessions for each monkey (Monkey A: F_2,8_ = 11.1, p = 0.004; Monkey O: F_2,12_ = 17.23, p = 0.0002; Figure 4A-B, E-F and left inset bar graphs). Tukey post-hoc tests showed a reduction of shoulder muscle activity from baseline to late adaptation phases (Monkey A: 44%, p = 0.002; Monkey O: 33%, p < 0.0001), and a quick return, becoming indistinguishable from the activity in baseline trials, when releasing the shoulder joint again in the post-adaptation phase (Monkey A: p = 0.483; Monkey O: p = 0.275). We found no corresponding changes in elbow TR muscle activity as a function of phases (one-way-ANOVA, Monkey A: F_2,8_ = 1.49, p = 0.28; Monkey O: F_2,12_ = 0.523, p = 0.605; Figure 4C-D, G-H and insets).

Moreover, we performed a t-test on the difference in shoulder muscle activity (adapt-baseline) in sessions in which the shoulder joint was fixed and was never fixed (control), and that monkeys performed a similar number of trials. Indeed, the reduction in shoulder muscle activity in sessions with the shoulder fixed was reliably smaller than at the equivalent point in the control sessions (Monkey A: t_6.5_ = -3.9, p = 0.005; Monkey O: t_6.8_ = -3.7, p = 0.007; Figure 4A, E and right inset bar graphs), indicating that shoulder muscle activity decay is related to shoulder fixation manipulation rather than the production of an large number of movements.

Additional evidence that monkeys adapted to the imposed novel arm dynamics with shoulder fixation was the presence of robust kinematic after-effects. That is, early in the post-adaptation phase, monkeys also generated substantial kinematic reaching errors, and those errors were in the direction expected (i.e., outwards of a reach arc of a pure elbow extension movement) when failing to compensate for intersegmental arm dynamics during elbow extension trials (Figure 5A-B). Note that this early release of the shoulder joint resulted in monkeys not achieving the speed and accuracy constraints imposed in our task. A one-way ANOVA comparing averaged reach errors (measured as hand distance from the center of the goal target at 80% of the movement, see Material and Methods) in trials late in the baseline (last 20 trials), early in the post-adaptation (first 3 trials) and late in the post-adaptation (last 20 trials) phases revealed a reliable effect of phase across sessions for each monkey (Monkey A: F_2,8_ = 12.9, p = 0.003; Monkey O: F_2,10_ = 8.26, p = 0.007; Figure 5A-B, bottom row). We chose a smaller bin size early in the post-adaptation because the return to baseline after unlocking the shoulder joint happens quickly (Figure 4A,E). Tukey post-hoc tests showed a reliable increase in kinematic errors from baseline to early-post-adaptation phases (Monkey A: 80%, p = 0.001; Monkey O: 81%, p < 0.0001), and a return to baseline accuracy levels late in the post-adaptation phase (Monkey A: p = 0.147; Monkey O: p = 0.025).

**Figure 5:**
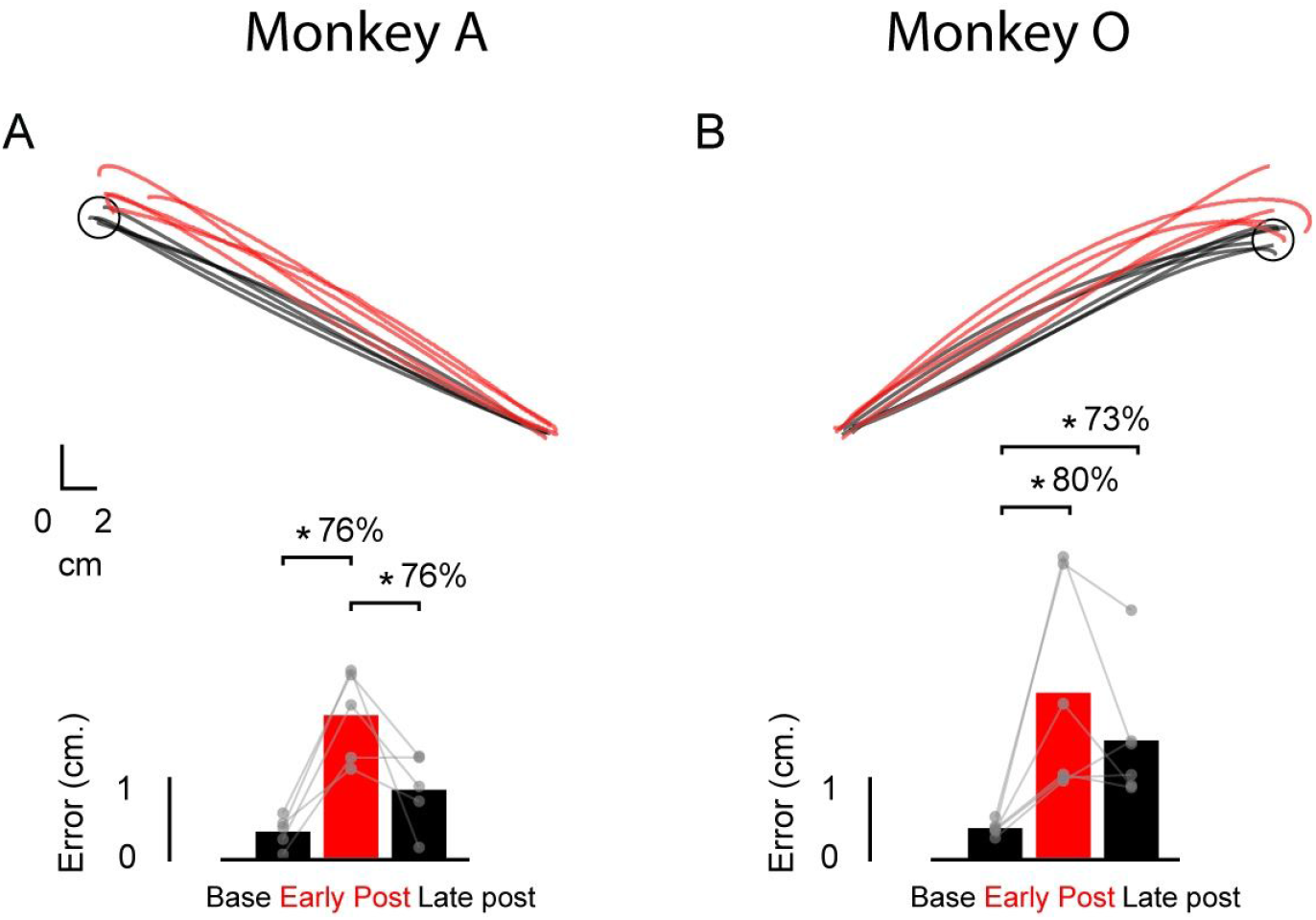
Hand trajectories before and after adapting elbow reaches with shoulder fixation. **A**, Average hand trajectories (top row) late in the baseline (20 trials) and early in the post-adaptation trials (first 3 trials) both with the shoulder joint free to rotate. Each trace represents data from a single session. Average error to the center of the target in the last 20 trials in the baseline, first 3 trials in the post-adaptation and last 20 trials in the post-adaptation. Each dot represents data from a single session. Asterisks indicate reliable effects (p < 0.05; see main text). **B**, Data for monkey O are shown using the same format as for monkey A in **A**.

### Transfer to feedback control

Our second main question was whether learning novel arm dynamics during feedforward reaching with shoulder fixation in monkeys also modifies feedback responses to mechanical perturbations.

Figure 6A, C left panel illustrates the average of the difference (excitatory - inhibitory conditions) of shoulder (PD) muscle activity in the long-latency epoch over trials late (last 3 trials) in the baseline, adaptation and post-adaptation phases of the protocol. Each dot represents the average EMG from a single session. A one-way ANOVA on this data revealed a reliable effect of phase on shoulder long-latency muscle responses across sessions for each monkey (Monkey A: F_2,8_ = 37.73, p < 0.0001; Monkey O: F_2,10_ = 8.04, p = 0.008). Tukey post-hoc tests showed a reliable reduction of shoulder long-latency responses from baseline to late adaptation phases (Monkey A: 23%, p = 0.001; Monkey O: 30%, p = 0.03), and a quick return to that seen in baseline levels (Monkey A: p = 0.01; Monkey O: p = 0.28).

**Figure 6:**
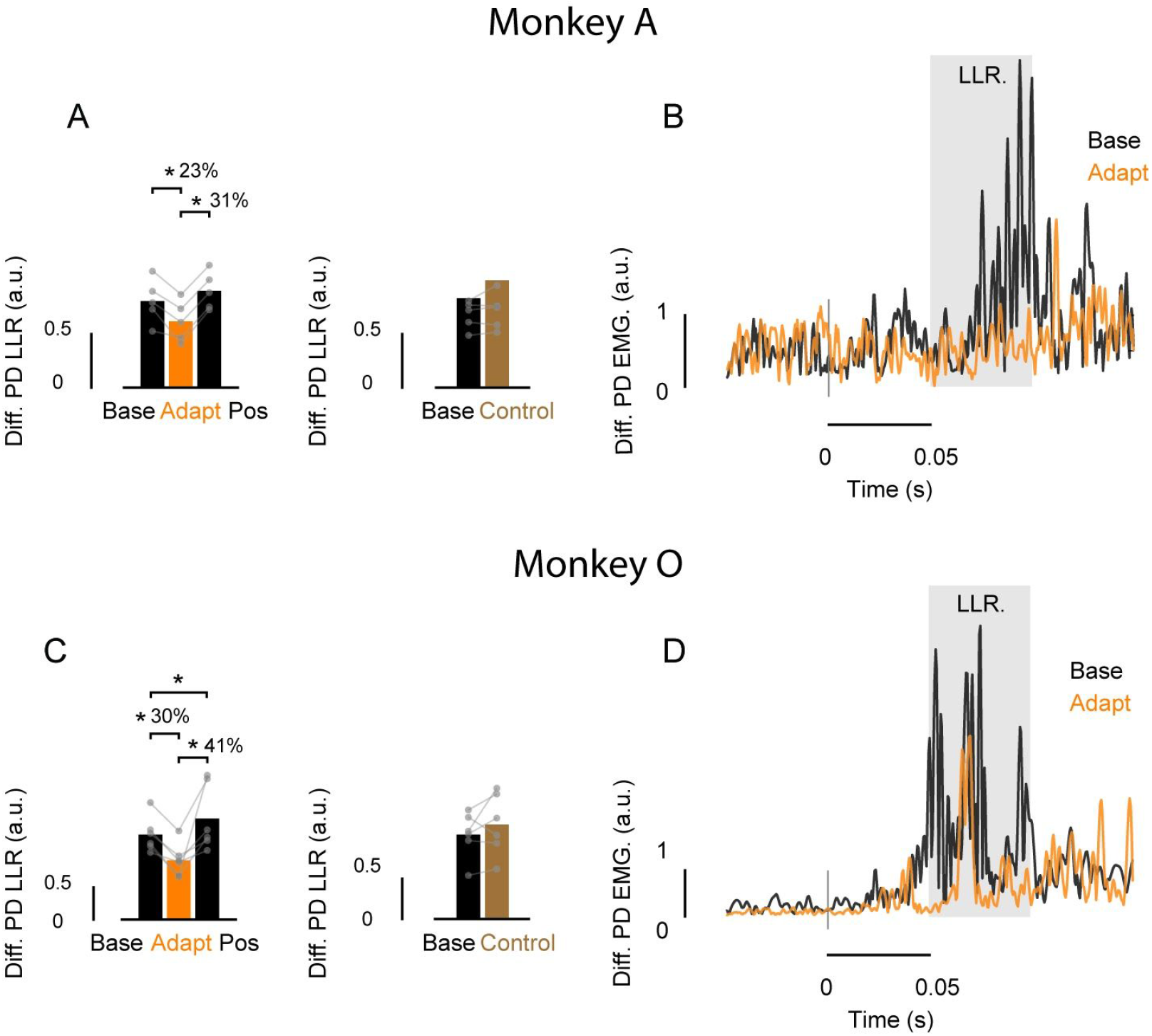
Rapid feedback responses before and after adapting self-initiated elbow reaching with shoulder fixation. **A**, Left bar graph shows the average of the difference (excitatory-inhibitory) of shoulder PD EMG (filtered, rectified and normalized) in the long latency epoch (50-100 ms) across trials late in the baseline, adaptation and post-adaptation (last 3 trials). Each dot represents data from a single session. Asterisks indicate reliable effects (p < 0.05; see main text). Right bar graph shows the data of the average of the difference (excitatory-inhibitory) of shoulder PD EMG in the long latency epoch across control sessions (first and last 3 trials) in which the shoulder joint was never fixed. **B**, Time series of normalized PD muscle activity in one exemplar session averaged over the last 3 baseline (black) and adaptation (orange) trials. Data are aligned on perturbation onset. Grey horizontal shaded area indicates the Long-Latency epoch (LLR). **C-D**, Data for monkey O are shown using the same format as for monkey A in **A-B**.

We found no reliable differences in EMG activity in a pre-perturbation epoch across learning phases (one-way-ANOVA, Monkey A: F_2,8_ = 9.32, p = 0.43; Monkey O: F_2,10_ = 9.97, p = 0.409, Figure 6B, D) suggesting that changes in the long-latency epoch do not merely reflect changes in the state of the motor neuron pool (gain scaling, Marsden *et al*., 1976; Bedingham & Tatton, 1984; Pruszynski *et al*., 2009). However, as we did not ensure that the muscle was active prior to perturbation onset, we cannot rule out this possibility (see limitation section below). We also found no corresponding changes in shoulder PD muscle response in the short-latency epoch as a function of the phases of the protocol (one-way-ANOVA, Monkey A: F_2,8_ = 2.442, p = 0.149; Monkey O: F_2,10_ = 0.07, p = 0.9, Figure 6B, D). Note that we used a combination of loads that created minimal shoulder joint motion, and thus it was expected minimal to no activity in this epoch even in baseline trials (Kurtzer *et al*., 2008, 2009; Maeda *et al*., 2018; Maeda, Zdybal, *et* al., 2020a). Moreover, we found no corresponding changes in elbow muscle responses in long-latency epoch as a function of experimental phase (one-way-ANOVA, Monkey A: F_2,8_ = 1.9, p = 0.21; Monkey O: F_2,10_ = 0.17, p = 0.84).

Importantly, we also extracted data from control sessions, where monkeys performed similar number of trials but never with the shoulder fixed, and we found no decrease in shoulder PD activity in the long-latency epoch early and late in these control sessions (paired t-test, Monkey A: t_7.7_ = -0.49, p = 0.63; Monkey O: t_9.2_ = -0.64, p = 0.536, Figure 6A, C, right panel).

## Discussion

Motivated by our previous work in humans (Maeda *et al*., 2018; Maeda, Gribble, *et al*., 2020; Maeda, Zdybal, *et al*., 2020a, 2020b), we first tested whether macaque monkeys learn to reduce shoulder muscle activity when performing self-initiating pure elbow movements with their shoulder joint mechanically fixed. Consistent with such learning, we found a systematic reduction in this shoulder muscle activity over trials when the shoulder joint was fixed. After releasing the shoulder joint again, we found evidence of kinematic aftereffects (i.e., reach errors), in the direction predicted if the animal was failing to compensate for the arm’s normal dynamics. We also tested whether such learning transfers to feedback responses to mechanical perturbation. Consistent with transfer, we found a reduction in shoulder long-latency feedback responses, as appropriate for the altered arm intersegmental dynamics. Together, our results demonstrate that macaque monkeys, like humans, learn novel intersegmental arm dynamics during self-initiated reaching with shoulder fixation, and that such learning transfers to feedback control (Maeda *et al*., 2018; Maeda, Gribble, *et al*., 2020; Maeda, Zdybal, *et al*., 2020a).

### Relationship to human studies

Our monkey tasks were chosen in a way to mimic several aspects of previous human studies. First, research has demonstrated that humans generate predictive shoulder muscle activity when generating pure elbow movements (Gribble & Ostry, 1999; Debicki & Gribble, 2005; Maeda *et al*., 2017, 2018) and show robust shoulder long-latency reflex responses to perturbations applied to the arm that creates pure elbow motion (Kurtzer, Pruszynski, *et al*., 2006; Kurtzer *et al*., 2008, 2009, 2014, 2016; Crevecoeur *et al*., 2012; Maeda *et* al., 2017, 2018; Kurtzer, 2019; Maeda, Gribble, *et al*., 2020). Likewise, when monkeys generate elbow reaches or respond to perturbations that created pure elbow motion in our study, shoulder muscle activity begins before movement onset (Figure 2C-D), and shows a quick shoulder responses within 50-100 ms post perturbation (i.e., long-latency epoch; Figure 3C-D), respectively. These results are consistent with previous work showing that monkeys generate muscle activity in a way that mirrors the control of the arm’s dynamics during multi-joint reaching (Scott, 1997; Graham *et al*., 2003; Kurtzer, Herter, *et al*., 2006) and perturbation tasks (Pruszynski *et al*., 2011).

Previous studies have shown that human participants learn to reduce shoulder muscle activity with shoulder fixation but such learning is slow (i.e., it unfolds over hundreds of trials) and is incomplete (i.e., activity is not eliminated) (Maeda *et al*., 2018; Maeda, Zdybal, *et al*., 2020a, 2020b). Although the rate of learning seems faster for monkey O than monkey A (Figure 4E) and indeed both monkeys showed faster learning than the typical human participant, learning was still incomplete for both monkeys in our study. The main difference between the current study with respect to human studies is that the animals repeat this learning protocol for many testing sessions. Thus, there could be an effect of savings across sessions (Krakauer, 2009), which, to our knowledge, has not yet been examined in human studies (see also the Limitations section below). After adapting to the novel arm’s dynamics, the monkeys show that rapid feedback responses are also adapted in a way consistent with the new underlying joint torques, also similar to humans in this task (Maeda *et al*., 2018; Maeda, Gribble, *et al*., 2020; Maeda, Zdybal, *et al*., 2020a). Taken together, the similarities at the behavioural and muscle activity levels suggest that these animals are sufficient for investigating the shared neural mechanisms for feedforward and feedback control of reaching.

### Learning novel for arm dynamics during self-initiated reaching

Like humans, monkeys can learn a variety of upper-limb motor tasks from reaching in the presence of force environments (Gandolfo *et al*., 2000; Li *et al*., 2001; Gribble & Scott, 2002; Arce *et al*., 2010; Richardson *et* al., 2012; Cherian *et al*., 2013; Perich *et al*., 2018) to visuomotor mapping representations (Paz *et al*., 2003; Mandelblat-Cerf *et al*., 2009; Perich *et al*., 2018; Vyas *et al*., 2018, 2020) and brain machine interface (i.e., BMI) control (Sadtler *etal*., 2014; Golub *et al*., 2015, 2018; Hennig *et al*., 2018; Vyas *et al*., 2018; Oby *et al*., 2019; Zhou *et al*., 2019). These types of learning, however, unfold as a function of errors the animals make in the task and are consistent with the monkey updating an internal model of the environment (for review on error-based learning see, Shadmehr *et al*., 2010). In our task, fixating the shoulder joint clamps reach trajectories to an arc rotating about the elbow joint and does not introduce large kinematic errors during learning. Reducing shoulder muscle activity in this task is efficient because fixing the shoulder joint cancels the forces that arise at the shoulder with forearm rotation, and thus removes the need to generate shoulder muscle activity. The observation that our monkeys show learning in this task in a way similar to humans expands the scope of the types of motor learning paradigms for which macaque monkeys can be a useful model (Maeda *et al*., 2018; Maeda, Zdybal, *et al*., 2020a, 2020b).

### Transfer to feedback control

Like humans, when monkeys learn new intersegmental dynamics during self-initiated movement such learning appears to transfer to feedback control in the context of mechanical perturbations. Current theories in sensorimotor neuroscience, based on optimal feedback control, posit that voluntary motor behavior involves the sophisticated manipulation of sensory feedback (Todorov & Jordan, 2002; Scott, 2004; Pruszynski & Scott, 2012). Consistent with this idea, a number of studies in humans have shown that learning new feedforward coordination patterns (like when reaching in a novel force field) automatically transfer to transcortical feedback responses (Wang *et al*., 2001; Kimura *et al*., 2006; Wagner & Smith, 2008; Kimura & Gomi, 2009; Ahmadi-Pajouh *et al*., 2012; Yousif & Diedrichsen, 2012; Cluff & Scott, 2013; Maeda *et al*., 2018; Maeda, Gribble, *et al*., 2020; Maeda, Zdybal, *et al*., 2020a). For instance, when human participants learn to reach in the presence of an external force environment or learn to reduce shoulder muscle activity when generating elbow reaching movements with shoulder fixation, evoked long-latency feedback responses to mechanical perturbations mirrors such learning during reaching. Here we found that such transfer is also present in monkeys. After monkeys learn novel arm dynamics during reaching with shoulder fixation, shoulder long-latency reflex responses to perturbations were also reduced.

This transfer between feedforward and feedback control is thought to take place because of shared neural mechanisms between feedforward and feedback control. One potential node that could mediate such a transfer system is the primary motor cortex (M1). Previous work in humans and monkeys has demonstrated that M1 mediates feedforward and feedback compensation for the arm’s dynamics (Gritsenko *et al*., 2011; Pruszynski *et al*., 2011), it mediates muscle coordination patterns (Morrow *et al*., 2009; Cherian *et al*., 2013; Sussillo *et al*., 2015; Russo *et al*., 2018; Naufel *et al*., 2019), and show single unit neural activity for both feedforward and feedback control (Evarts, 1973; Evarts & Tanji, 1976; Wolpaw, 1980; Evarts & Fromm, 1981; Picard & Smith, 1992; Herter *et al*., 2009; Pruszynski *et al*., 2011; Omrani *et al*., 2014; Pruszynski, 2014). M1 is likely not mediating such motor learning and transfer alone (Cherian *et al*., 2013; Perich *et al*., 2018). For example, M1 is highly connected with the cerebellum (Allen & Tsukahara, 1974; Hoover & Strick, 1999; Holdefer *et al*., 2000; Wagner *et al*., 2019), which is also a site that has been long proposed to contain the computations for internal models that underlie feedforward motor control (Wolpert & Kawato, 1998; Wolpert *et al*., 1998). The cerebellum is also critical for feedforward and feedback control of the intersegmental arm dynamics, as a deficit in this region in patients has been reported to substantially impact the ability to coordinate across joints (Holmes, 1939; Goodkin *et al*., 1993; Bastian *et al*., 1996, 2000; Kurtzer *et al*., 2013). Demonstrating learning and transfer in monkeys, as we do here, sets the stage for further neurophysiology experiments for hunting these potential neural mechanisms of feedforward and feedback control of the arm dynamics.

### Limitations

As always, there are limitations to this study that need to be explicitly noted. First, the monkeys performed the perturbation tasks without a pre-perturbation background load. Although we found no changes in the levels of muscle activity in the pre-perturbation epoch, the lack of background load means that it is not possible to rule out small changes in the state of the motor neuron pool as a function of learning (gain scaling, Marsden *et al*., 1976; Bedingham & Tatton, 1984; Pruszynski *et al*., 2009) explaining the transfer between feedforward and feedback control. The reason for not using background loads is because it reduces effort for the monkey and thus training burden. Moreover, because the monkeys were being used for cortical neural recordings, such background loads were less critical. Second, as already described above, the learning patterns appeared somewhat different across the two animals with monkey A showing slow and gradual learning similar (though still faster) to humans whereas monkey O showed a more immediate reduction in shoulder muscle activity after shoulder fixation. We do not have any definitive explanation for these differences or how they relate to the underlying neural mechanisms though the monkeys had more experience with the task than a typical human participant and, in general, individual humans also show some variability in learning rates (Krakauer *et al*., 2019). Third, note that similar to the human studies that investigated these same behavioural and EMG metrics (Maeda *et al*., 2018; Maeda, Zdybal, *et al*., 2020a), it is not possible to casually link the reduction of the feedforward motor commands with the reduction of the long-latency reflex responses without further invasive methods and neural recordings.

## Acknowledgements

We thank Rhonda Kerstern and Whitney Froese for their technical support.

